# Novel Insights into Selection for Antibiotic Resistance in Complex Microbial Communities

**DOI:** 10.1101/323634

**Authors:** Aimee K. Murray, Lihong Zhang, Xiaole Yin, Tong Zhang, Angus Buckling, Jason Snape, William H. Gaze

**Affiliations:** Medical School, University of Exeter, Environment & Sustainability Institute, Penryn Campus, Cornwall, TR10 9FE; Department for Civil Engineering, University of Hong Kong, Pokfulam, Hong Kong; College of Life and Environmental Sciences, University of Exeter, Daphne Du Maurier Building, Penryn Campus, Cornwall, TR10 9FE; AstraZeneca Global Environment, Mereside, Alderley Park, Macclesfield, Cheshire, SK10 4TG

## Abstract

Recent research has demonstrated selection for antibiotic resistance occurs at very low antibiotic concentrations in single species experiments, but the relevance of these findings when species are embedded in complex microbial communities is unclear. We show the strength of selection for naturally occurring resistance alleles in a complex community remains constant from low sub-inhibitory to above clinically relevant concentrations. Selection increases with antibiotic concentration before reaching a plateau where selection remains constant over a two order magnitude concentration range. This is likely to be due to cross-protection of the susceptible bacteria in the community following rapid extracellular antibiotic degradation by the resistant population, shown experimentally through a combination of chemical quantification and bacterial growth experiments. Metagenome and 16S rRNA analyses on sewage-derived bacterial communities evolved under cefotaxime exposure show preferential enrichment for *bla*_CTX-M_ genes over all other beta-lactamase genes, as well as positive selection and co-selection for antibiotic resistant, opportunistic pathogens. These findings have far reaching implications for our understanding of the evolution of antibiotic resistance, by challenging the long-standing assumption that selection occurs in a dose-dependent manner.

## IMPORTANCE

Antibiotic resistance is one of the greatest, global issues facing modern society. Still, comparatively little is known about selection for resistance at very low antibiotic concentrations. We show that the strength of selection for clinically important resistance genes within a complex bacterial community can remain constant across a large antibiotic concentration (wide selective space). Therefore, largely understudied ecological compartments could be just as important as clinical environments for selection of antibiotic resistance.

## INTRODUCTION

Antibiotic resistance poses a major threat to society, the sustainability of modern healthcare systems, food security and the global economy [1, 2]. Until recently, most research on evolution of resistance focused on selection at clinically relevant antibiotic concentrations, as the ‘traditional’ selective window hypothesis was universally accepted. This hypothesis states that selection for antibiotic resistance will only occur above the minimum inhibitory concentration (MIC) of susceptible bacteria and below the MIC of resistant bacteria [3]. In fact, numerous experimental studies have observed selection for resistance at sub-MIC antibiotic concentrations, at the point where the selective pressure (antibiotic) is sufficient to offset the cost of resistance [3–7]. In recent isogenic studies, a single host species with chromosomal or plasmid-borne resistance mechanisms were competed with their susceptible counterparts at varying concentrations of antibiotic to determine the minimal selective concentration (MSC) [3, 5]. The MSC is the lowest concentration of antibiotic at which resistance is positively selected, which can be significantly lower than the minimum inhibitory concentration (MIC) [3, 5]. MSCs have also been estimated using publically available, clinical breakpoint data [8] but experimental data is required to assess the validity of these predictions, especially in a community context. These findings show that the selective compartment (meaning the antibiotic gradient and spatial range along which resistant bacteria/genes could be enriched) is much larger than previously thought [9]. This in turn suggests selection may be occurring in previously unconsidered selective compartments which harbour relatively low antibiotic concentrations, such as the gut microbiome, waste water and even surface waters contaminated with antibiotic residues.

Though these findings are significant, the use of single species means their relevance with regards to selection in complex microbial communities remains unclear. Many studies have quantified numbers and/or prevalence of resistance genes in waste water influent and effluent (including, but not limited to [10, 11]), with a recent study utilising epicPCR to identify the host background of highly abundant resistance genes [12]. However, positive selection for resistance within complex bacterial communities is a current knowledge gap [13], with experimental MSC data in complex communities severely lacking. One recent study [14] reported a biological effect at low concentrations of tetracycline in a microbial community, by quantifying tetracycline resistance gene prevalence (*tetA* and *tetG* genes normalised to 16s rRNA copy number). However, as starting gene frequencies were not measured, it is unclear if the observed effect was driven by positive selection; or reduced negative selection. In other words, without comparing the final resistance gene prevalence to the initial resistance gene prevalence, it is unknown if resistance genes actually increased over time under tetracycline exposure (i.e. were positively selected); or if resistance genes were simply lost at a slower rate compared to the no antibiotic control (i.e. were negatively selected, or showed increased persistence). Here, we aimed to quantify positive selection in a complex bacterial community by conducting evolution experiments using a waste water bacterial community inoculum to determine the MSC of cefotaxime. Co-selection for other resistance genes and effects on community structure were also determined through metagenome analyses.

Cefotaxime is a World Health Organisation (WHO) recognised ‘critically important’ antibiotic [15], ‘essential’ for human medicine [16] that was most recently identified as a key antimicrobial stewardship target through inclusion in the WHO ‘watch list’ of essential medicines [17]. In this study, prevalence of the *bla*_CTX-M_ gene group was determined with qPCR and selection coefficients were calculated to estimate the MSC of cefotaxime. CTX-Ms are extended spectrum beta-lactamases (ESBLs) which cleave the beta-lactam ring, effectively inactivating and degrading beta-lactam antibiotics [18]. Previous work has demonstrated beta-lactamases can inactivate extracellular beta-lactams, to the benefit of nearby susceptible bacteria [19–21]. This protective effect on susceptible bacteria has since been shown for an intracellularly expressed resistance mechanism, also degradative in nature [22].

Results show, for the first time, that selection for *bla*_CTX-M_ genes occurs at very low, sub-inhibitory concentrations. We also demonstrate that selection occurs with equal potency at very low antibiotic concentrations and at concentrations greatly exceeding those used in the clinic. Therefore, antibiotic resistance is not always selected for in a dose-dependent manner. These findings illustrate the importance of studying selection for resistance within complex bacterial communities over a wide selective range, representative of different selective compartments [9].

## RESULTS

**Cefotaxime exposure affects community structure.** Complex community (raw, untreated waste water) microcosms were spiked with a range of concentrations of cefotaxime. The exposure concentration range was selected from the EUCAST [23] defined clinical breakpoint concentration (the concentration at which *Enterobacteriaceae* are considered ‘clinically’ resistant) down to 0, in a two-fold dilution series including concentrations similar to those previously measured in different environments (such as hospital effluent, waste water treatment plant effluent and surface waters (low μg/L up to 150 μg/L [24–26])). Bacterial communities were transferred daily into fresh medium and fresh antibiotic for 8 days. Chemical quantification was performed at the beginning of day 0 and after 24 hours, to determine an accurate MSC (based on measured rather than nominal antibiotic concentration) and to assess the chemical stability of cefotaxime (a third generation cephalosporin of the beta-lactam class of antibiotics) in the system.

At the end of the experiment, three replicates from a low, medium and high antibiotic concentration were selected to undergo metagenome analyses, alongside the unexposed control to identify the key resistance genes under selection. Metagenomic DNA was sequenced with Illumina MiSeq2.

16S rRNA data was extracted from trimmed, quality controlled paired reads with MetaPhlan2 [27]. Overall, the community comprised of predominantly Gram negative bacteria, though Gram positive bacteria were also detected (Figure S1). Between-sample variation was expected due to unavoidable heterogeneity within the complex community inoculum. Even so, there were clear differences between the control untreated bacterial community and the communities exposed to cefotaxime amongst the 25 most abundance species (Figure 1, all detected species can be seen in Figure S1). In particular, several species were eliminated by cefotaxime treatment (or reduced below the limit of detection). These included the opportunistic Gram negative pathogens *Providencia alcalifaciens, Aeromonas veronii, Morganella morganii* and *Klebsiella pneumoniae*; as well as the opportunistic Gram positive pathogen *Streptococcus infantarius*. All of these were significantly associated with the no antibiotic control as determined by Linear Discriminant Analyses (LDA) effect size (LEfSe) analyses (Figure S2). Conversely, several Gram negative and Gram positive opportunistic pathogens showed greater abundance in treated communities compared to the untreated control; namely *Pseudomonas aeruginosa, Acinetobacter baumanii*, *Bacteroides fragilis* and *Enterococcus faecalis*; however, only *P. aeruginosa* was significantly enriched in the 2 mg/L cefotaxime treatment (LEfSe, Figure S2). Cefotaxime treatment also resulted in slightly decreased numbers of *Escherichia coli*, though this was not significant and it was still the predominant species across all treatments. In general, there was much greater variability between treatment replicates compared to the untreated control (Figure S1).

**Figure 1.**
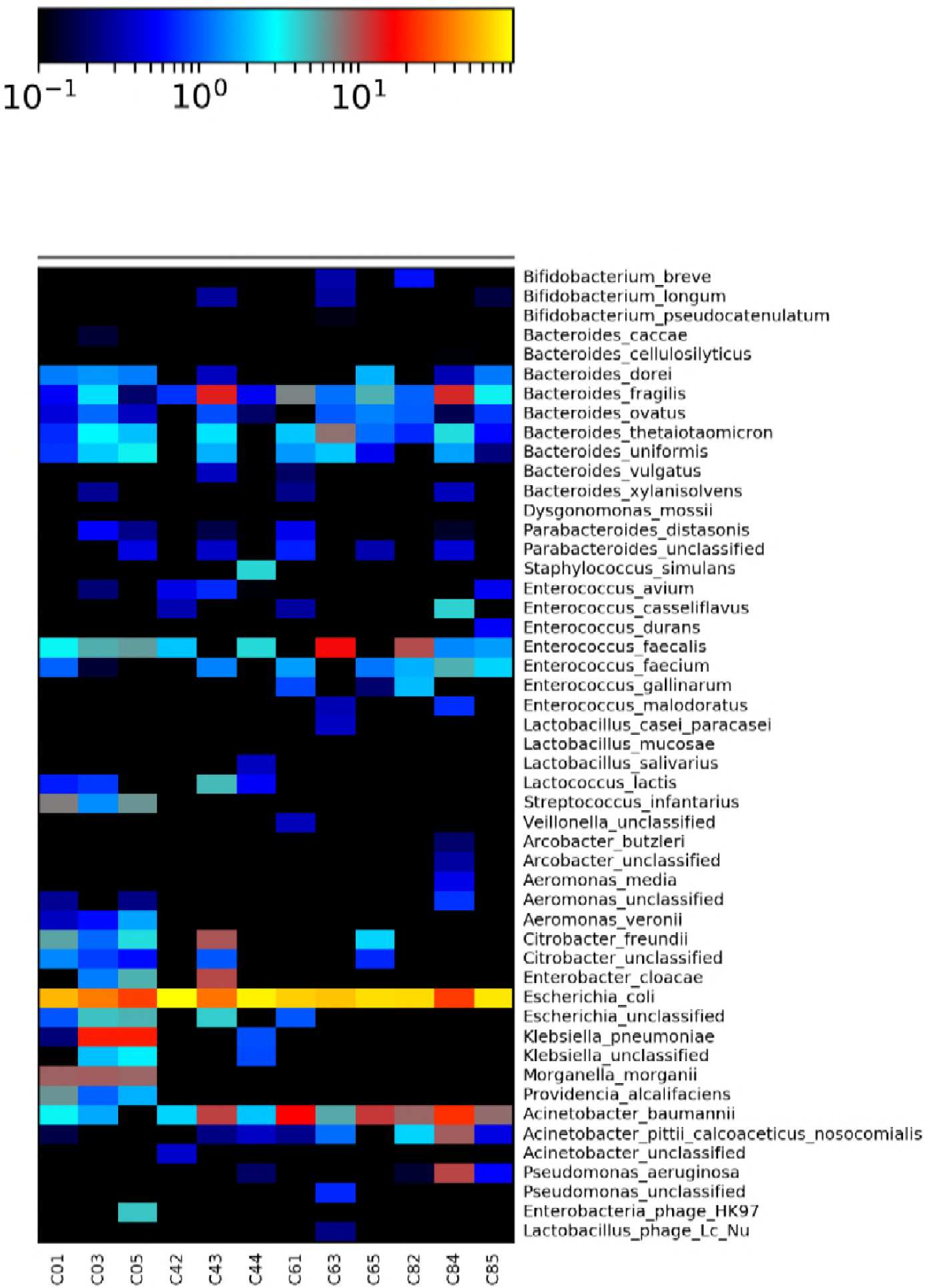
Heatmap showing the relative abundance of detected species using Bray Curtis distance measurements for treatment (x axis) and species (y axis) for each cefotaxime treatment. ‘C0’, ‘C4’, ‘C6’ and ‘C8’ correspond to 0, 125, 500 and 2000 μg/L cefotaxime respectively. The number after the concentration denotes the biological replicate number (1 − 5), chosen randomly for sequencing at day 8 of the experiment.

***Bla*_CTX-M_ genes are preferentially selected over all other beta-lactam resistance mechanisms.** Metagenomic data was further analysed with the ARGs-OAP pipeline [28], designed to thoroughly interrogate metagenomic data and identify resistance genes. Selection for beta-lactam resistance was prominent (as expected); however, co-selection for resistance to unrelated antibiotic classes was also observed: namely, co-selection for resistance to macrolides, aminoglycosides, trimethoprim, tetracyclines and sulphonamides which is likely to be due to carriage of multi-resistance plasmids (see Figure S3).

We delved deeper into the beta-lactam resistance genes to determine which genes, if any, were preferentially selected. We observed substantial enrichment for the beta-lactamase and extended-spectrum beta-lactamase (ESBL) genes *bla*_TEM_, *bla*_OXA_ and *bla*_CTX-M_ (Figure 2). Average increases in relative abundance from the lowest to highest concentration were 8-fold, 8-fold and 70-fold, respectively. *Bla*_CTX-M_ was preferentially selected over all other beta-lactamase encoding genes at each cefotaxime concentration.

**Figure 2.**
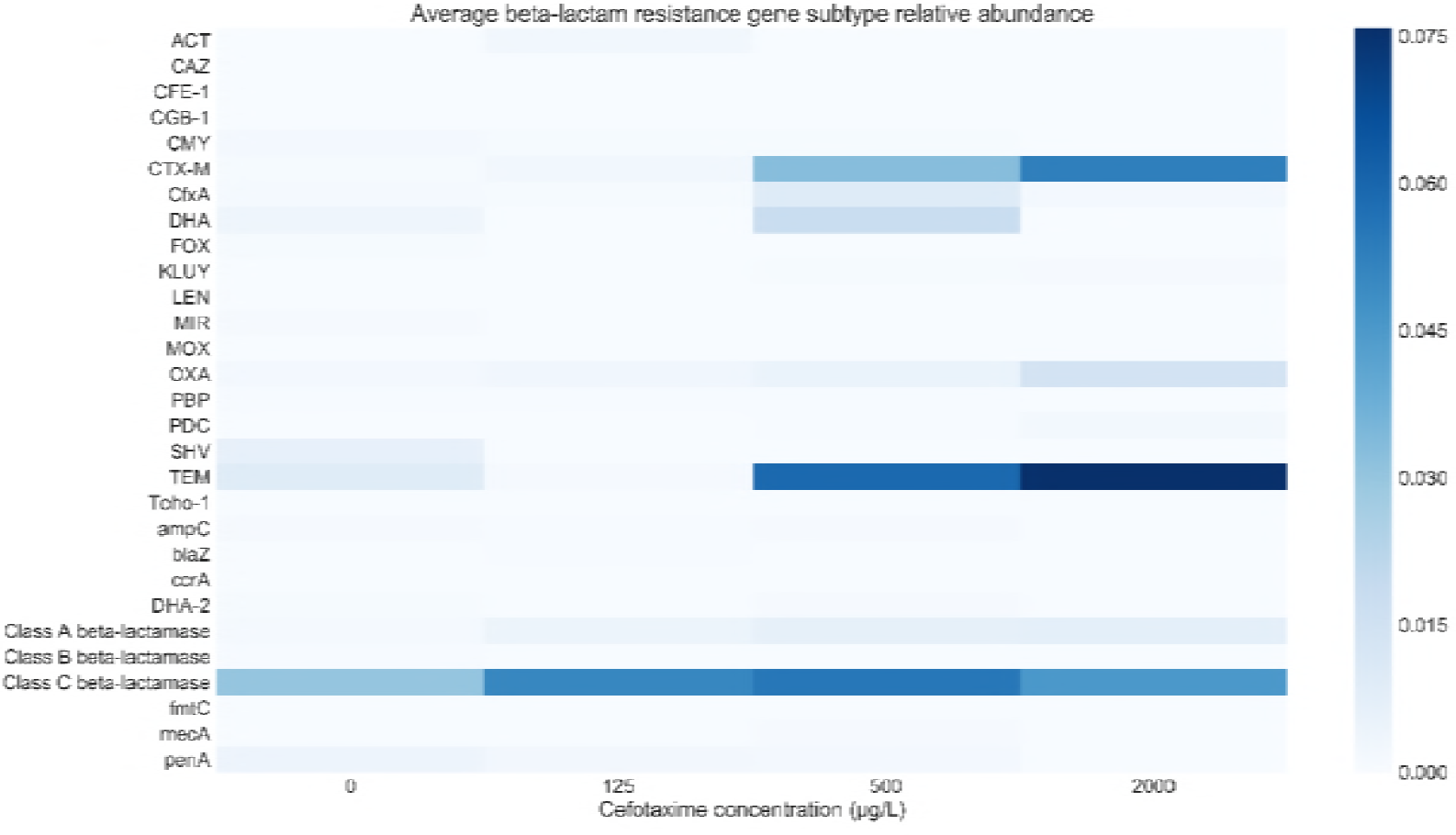
Heatmap showing average (n = 3) detected beta-lactam resistance gene subtype relative abundance (resistance gene number normalised with 16S rRNA copy number), following 8 days culture with cefotaxime. Only genes detected with the ARGs-OAP pipeline are shown.

**The MSC of cefotaxime is very low, but selection plateaus across a large concentration range.** Given the strong positive selection for *bla*_CTX-M_, we focused on accurate quantification of this group of genes across the entire experimental antibiotic gradient using qPCR. This follows previous work which showed qPCR is the most sensitive method for MSC determination [14]. *Bla*_CTX-M_ gene copy number was normalised to 16S rRNA copy number, to determine a molecular ‘prevalence’ of *bla*_CTX-M_; this prevalence was determined for each cefotaxime concentration both at the beginning and end of the experiment. A Kruskal-Wallis test confirmed *bla*_CTX-M_ prevalence, 16S rRNA copy number and *bla*_CTX-M_ copy number (all n=5 each) did not differ significantly between treatments at day 0.

Selection coefficients based on change in *bla*_CTX-M_ prevalence over time were calculated and plotted against cefotaxime concentration as in previous single species assays [3, 5] (Figure 3). A positive selection coefficient value indicates positive selection is occurring, and the x-axis intercept estimates the MSC; here, 0.4μg/L (Figure S4).

**Figure 3.**
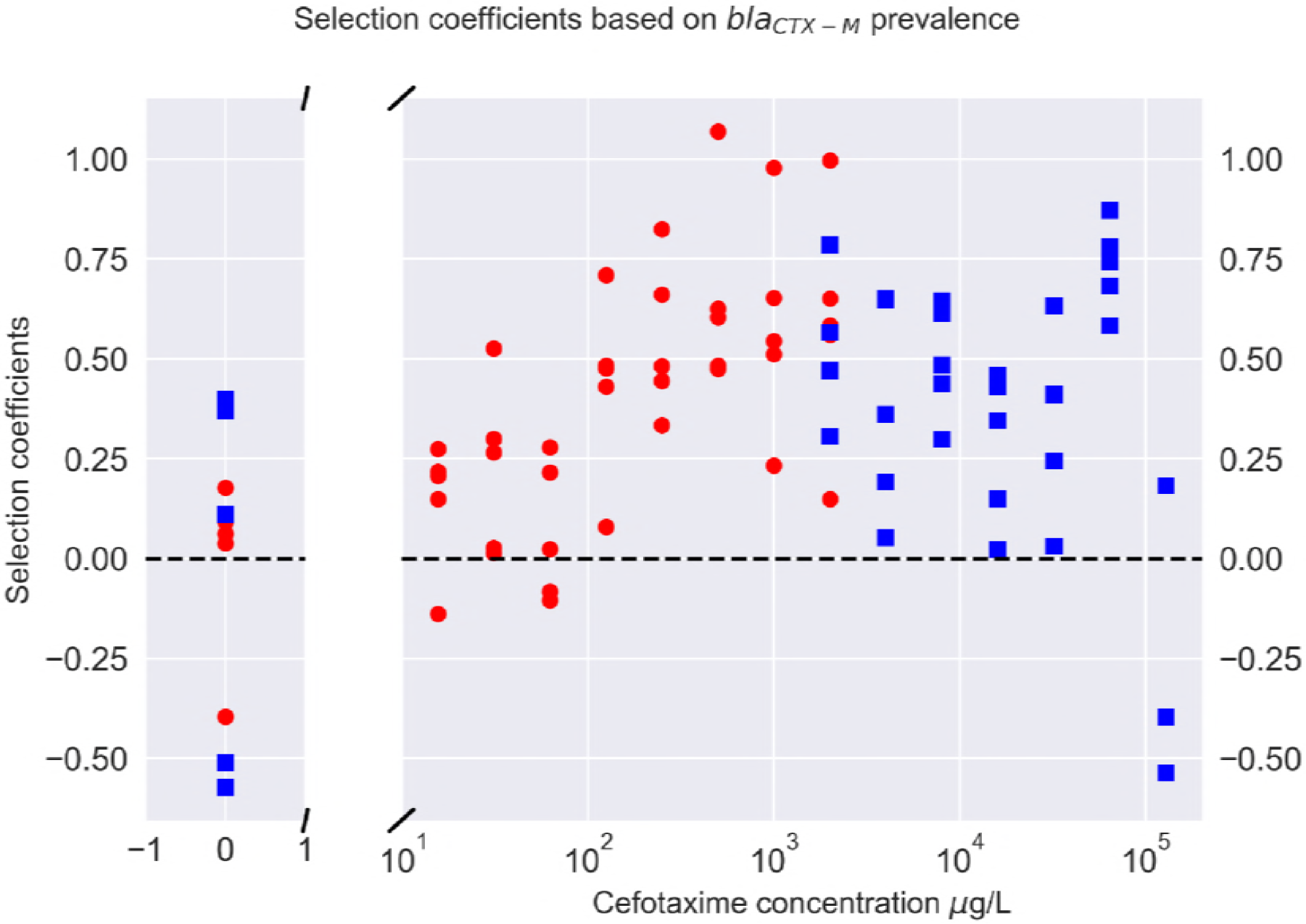
Selection coefficients (n=5) for each cefotaxime concentration, which equal the natural log of resistance gene prevalence (bla_CTX-M_ gene/16S rRNA copy number) at day 0 and day 8. 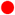=selection coefficients from low concentration experiment and 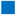 =from high concentration experiment. Selection coefficients > 0 indicate positive selection.

*Bla*_CTX-M_ prevalence increased over time (Figure S5) and with antibiotic concentration (linear term: *F*_1_, 42 = 26.3, *P*<0.001), but appeared to plateau at 500μg/L (quadratic term: *F*_1_, 42 = 13.2, *P* < 0.001) so an additional experiment was performed to determine if this plateau continued at higher concentrations (Figure 3). As hypothesised, *bla*_CTX-M_ prevalence increased when exposed to cefotaxime (linear term: *F*_1_, 36 = 9.6, *P* <0.01) but remained relatively constant (quadratic term: *F*_1_, 36 = 9.4, *P* <0.01) up until the two highest concentrations used in this study (Figure 3). These concentrations are over 30x and 50x times the defined clinical breakpoint cefotaxime concentration of 2 mg/L for *Enterobacteriaceae*. The rise in bla_CTX-M_ prevalence at 64 mg/L was due to an increase in bla_CTX-M_ gene copy number, and the decrease at 128 mg/L was due to a significant decrease in bla_CTX-M_ and slight reduction in 16S rRNA copy number (Figures S6 and S7).

**The bacterial community readily degrades cefotaxime.** We hypothesised that this plateau in selection was due to both the mechanism and sociality of the *bla*_CTX-M_ genes: as beta-lactamase enzymes can be found both intracellularly and extracellularly [21], the plateau in *bla*_CTX-M_ prevalence may be due to negative frequency-dependent selection [19]. In other words, the more prevalent *bla*_CTX-M_ becomes, the lower its fitness as cefotaxime degradation is accelerated to the benefit of the entire community, including non-*bla*_CTX-M_ bearing competitors. To investigate if cefotaxime was degraded by the community, chemical quantification of cefotaxime in the presence of the community was performed. Incubating the microcosms for 24 hours resulted in complete degradation of cefotaxime, at all but the highest concentration (Table S1). All measured concentrations were lower than expected, and the lowest concentration (15.625 μg/L) was below the limit of detection at the beginning of the assay. Therefore the MSC (0.4 μg/L) is estimated based on nominal concentrations, but in reality is likely to be lower still. As cefotaxime is known to be relatively unstable [29] an overnight degradation experiment was conducted to determine the amount of biotic and abiotic degradation occurring in the experimental system. A sterile microcosm and another inoculated with the complex community was incubated and destructively sampled at 0 hours, 6 hours and then every 3 hours for 24 hours. In sterile culture cefotaxime had only partially degraded over 24 hours; whereas in the presence of the community, cefotaxime was undetectable following 12 hours incubation (Figure 4A and 4B). This increase in degradation rate coincided with the beginning of the exponential growth phase of the community (Figure S8).

**Figure 4.**
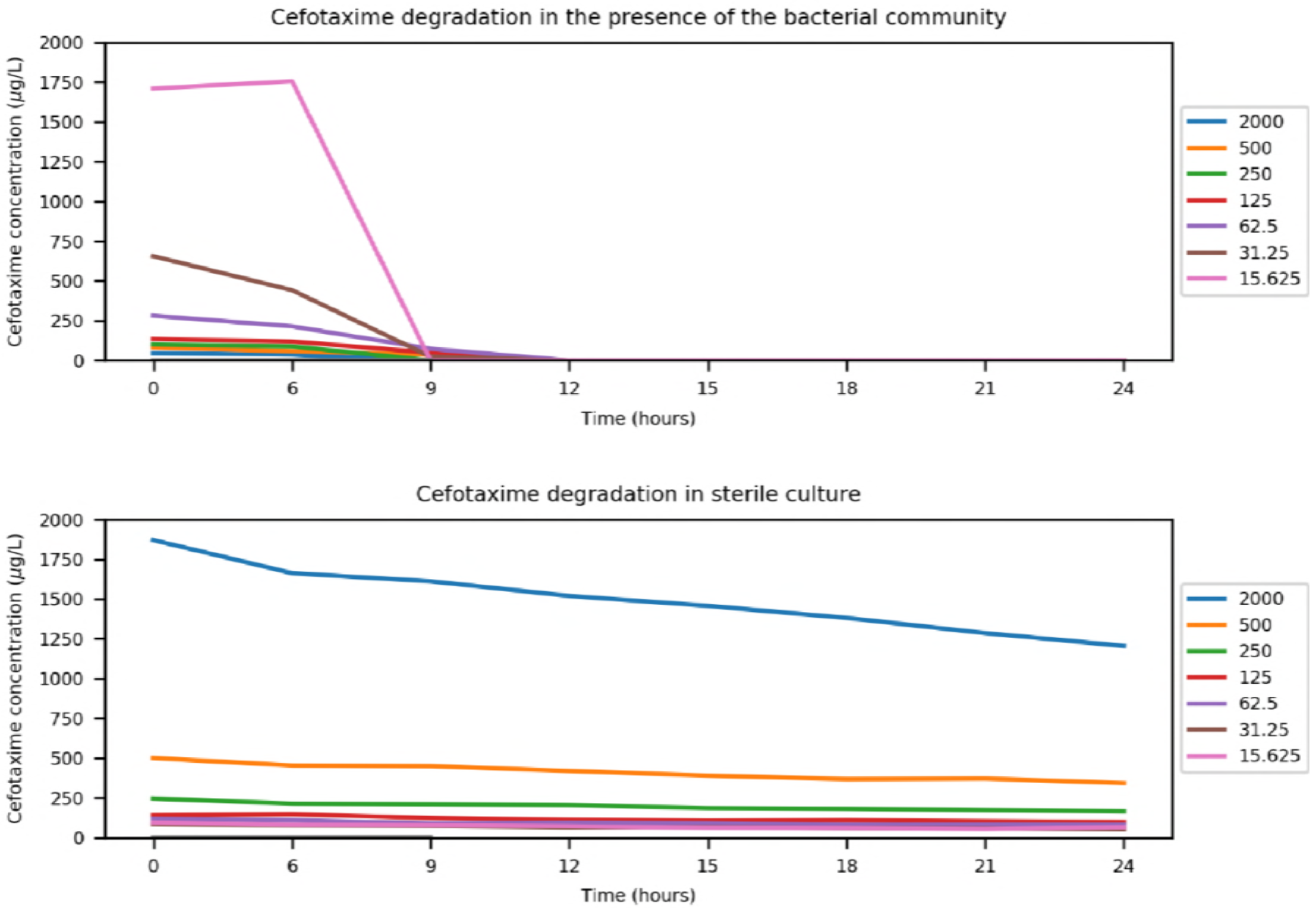
Single biological replicate, duplicate chemical replicate chemical quantification of cefotaxime (‘Measured cefotaxime concentration ug/L) at 0 and 6 hours, then every 3 hours for 24 hours at different starting cefotaxime concentrations (ug/L) in the presence of the complex bacterial community (A) and in sterile culture (B).

**Extracellular beta-lactamases ‘protect’ susceptible bacteria at cefotaxime concentrations well above MIC.** To confirm this degradation was biotic and by extracellular beta-lactamases, an additional experiment was performed whereby a susceptible *E. coli* strain (J53) was cultured in the presence of supernatant derived from an overnight culture of an *E. coli* strain bearing *bla*_CTX-M_-15, *bla_TEM-1_* and *bla*_OXA-1_ on a fully sequenced resistance plasmid [30] (strain NCTC 13451, available from Public Health England). Addition of the supernatant from the resistant strain allowed growth of the susceptible strain at the clinical breakpoint concentration [23], which was over 10x the MIC of the susceptible strain (Figure 5).

**Figure 5.**
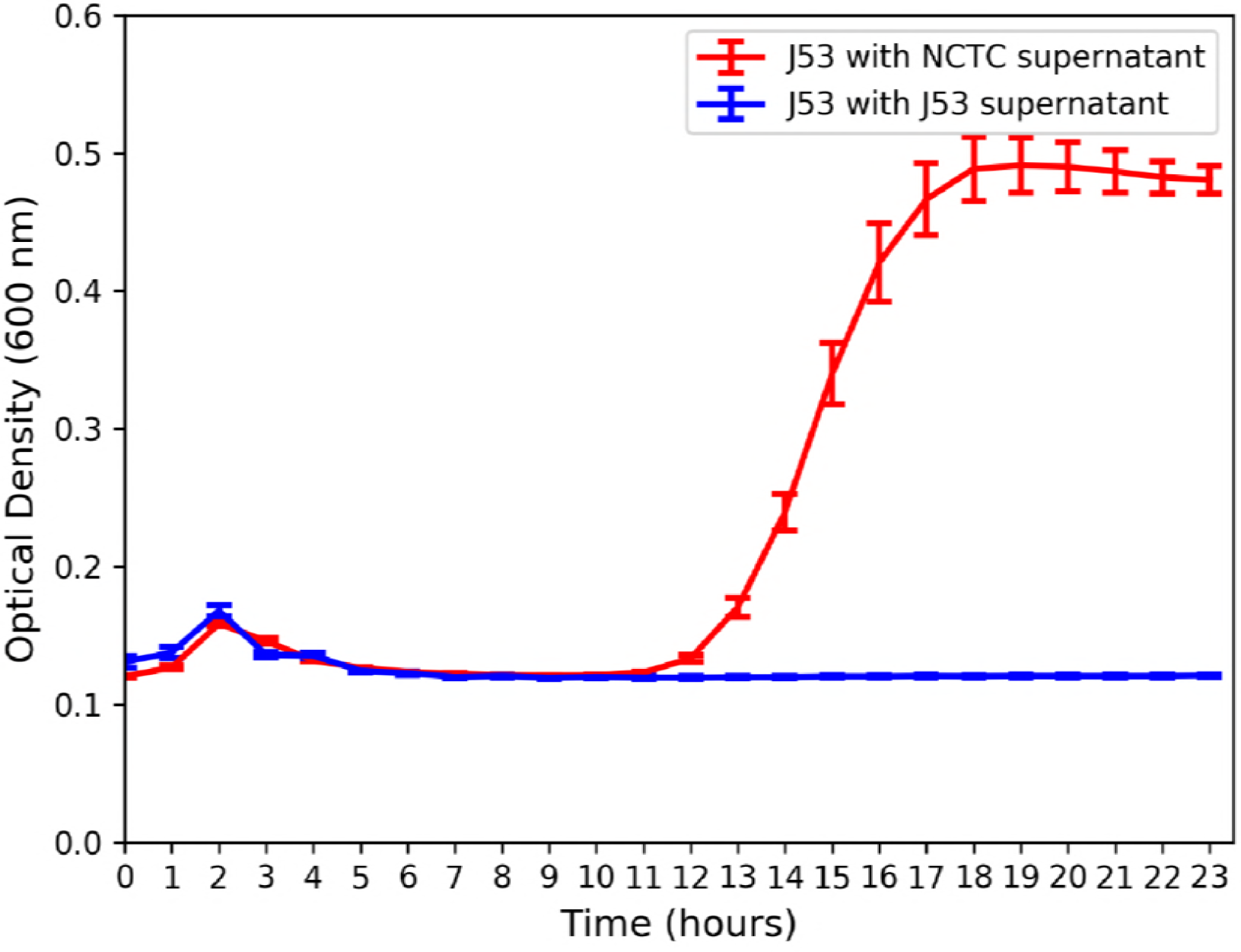
Average (n=4) optical density (600nm) over time of susceptible *E. coli* strain J53 grown in 2000 μg/L cefotaxime (clinical breakpoint concentration for *Enterobacteriaceae)* with beta-lactamase containing supernatant (NCTC strain 13451) or beta-lactamase free supernatant (strain J53).

## DISCUSSION

Here, we quantified positive selection for antibiotic (cefotaxime) resistance in a waste water derived complex bacterial community, by quantifying changes in resistance gene prevalence over time. We show clinically important [31] resistance genes (*bla*_CTX-M_) were positively selected at very low, environmentally-relevant concentrations likely due a combination of clonal expansion of hosts carrying *bla*_CTX-M_ and horizontal gene transfer of plasmids bearing *bla-*_CTX-M_. Antibiotic quantification has been identified as an overlooked aspect of MSC determination [13]. We quantified antibiotic concentrations when determining minimal selective concentrations and found cefotaxime to be rapidly degraded by the community, suggesting the estimated MSC of 0.4 μg/L to be an underestimate. Even so, the cefotaxime MSC determined in this study was very similar to several measured environmental concentrations [25, 32], suggesting selection could occur in certain environments such as hospital effluent and waste water influent. Responses to selection in such environments may be reduced, as unenriched bacterial communities e.g. in sewage may be impacted by high cell numbers and associated reduction in resource availability. However it is also possible that sustained exposure over long time periods would produce the same response.

In addition, we observed a plateau in the strength of selection across a very large antibiotic concentration range. This novel finding contrasts with previous work which has shown resistance to increase monotonically with antibiotic concentration [3, 5, 14]. A crucial implication of this finding is that selection for clinically important resistance mechanisms, such as *bla*_CTX-M_, may occur to a similar extent at sub-inhibitory concentrations as at high, clinical concentrations. The observed plateau in resistance selection has clinical relevance, when considering the antibiotic concentration gradients which inevitably form in different body compartments during chemotherapy [33]. These may also provide greater potential for selection for antibiotic resistance *in vivo* than previously considered. Potential overtreatment with unnecessarily long antibiotic courses [34] may compound this effect. Future research should address this finding and its relevance to environmental protection, effective antibiotic treatment and antimicrobial stewardship.

The observed plateau in selection for resistance is likely due to the cross-protective effect conferred by the resistant fraction on the susceptible fraction of bacteria in the population. Three lines of evidence strongly support this: 1) the degradative effect of the community, whereby within 24 hours all cefotaxime is degraded below the limit of chemical quantification, including the very highest, clinical breakpoint concentration. 2) metagenome analyses of 3 replicates at 4 antibiotic concentrations, which showed the main mechanism of resistance to the treatment antibiotic was degradative in nature, and could therefore provide a benefit to susceptible competitors within the community. 3) The single species *E. coli* experiment, which used supernatant from a resistant strain bearing a multi-resistance plasmid with the plasmid-free strain to show the potential extent of this community-wide benefit. The level of protection conferred by extracellular beta-lactamases in the supernatant of the resistant strain culture allowed growth of the susceptible strain well above its own MIC, at the clinical breakpoint concentration. Extrapolating this finding to the community, we hypothesise that within each 24 hour time period, CTX-M producers (and possibly other degraders) are selected for by cefotaxime, and are then outcompeted by susceptible bacteria following antibiotic degradation. This means resistant genotypes are likely to persist at even very low antibiotic concentrations, as they provide a benefit to the whole community; this effect has been modelled previously [19].

Selection for *bla*_CTX-M_ genes is likely due a combination of clonal expansion of hosts carrying *bla*_CTX-M_ and horizontal gene transfer of plasmids bearing *bla-*_CTX-M_. Our results are consistent with epidemiological data on beta-lactam resistance genes [35, 36], which document the rapid spread of *bla*_CTX-M_ genes worldwide to a ‘pandemic’ status. In this study, *bla*_CTX-M_ genes were under stronger selection than a large diversity of other resistance genes, possibly due to lower fitness cost (either metabolic or due to genetic context), and/or due to more efficient degradation and a potential wider degradative capacity. For example, previous research found the MICs of *bla*_CTX-M_ positive bacteria isolated from river sediment downstream of a waste water treatment plant were in excess of 2048 μg/ml [37]. Additionally, many TEM and OXA beta-lactamases do not have the extended-spectrum degradative capability of CTX-M ESBLs [35, 36]. We suggest the ability of *bla*_CTX-M_ genes to outcompete other beta-lactamase genes at all studied concentrations may also have contributed to the ‘pandemic’ spread of *bla*_CTX-M_ genes worldwide [35] and the replacement of other beta-lactamase variants [36].

The metagenome analyses showed *ampC* genes were detected but not enriched by cefotaxime exposure. Overexpression of chromosomal *ampC* genes can increase levels of resistance to many antibiotics, including cefotaxime but these genes were very rare within metagenomes and only confer low level resistance up to 8 mg/L [38] suggesting they do not play a significant role at the community level in this study [39]. The metagenome analyses also showed cefotaxime can also co-select for resistance to a range of antibiotic classes, even at sub-inhibitory cefotaxime concentrations. Co-occurrence of *bla*_CTX-M_ genes and aminoglycoside resistance has been reported previously and is confirmed here [36]. High levels of co-selection also indicates presence of multi-drug resistant plasmids, via co-resistance (i.e. co-localisation of multiple genes, of which only one needs to be under positive selection) [40].

The bacterial community analyses showed that cefotaxime had significant effects on community structure, even at sub-inhibitory concentrations through elimination of several species as determined by LEfSe. The bacterial communities in all treatments comprised mainly of Gram negative bacterial families, with *E. coli* being the most abundant species in all treatments. In waste water, with a lower temperature and different nutritional composition, inter-species competition with other taxa could result in reduced *E. coli* abundance which may be favoured by the media and temperature used in this study. However, *E. coli* are known to survive waste water treatment and are used as faecal indicator organisms to assess water quality [41]. Therefore, in waste water, *E. coli* may still possess a competitive advantage. There was much greater variability between replicates (for all species other than *E. coli*) within exposed communities, indicative of potential founder effects on evolution within individual microcosms, again supporting the hypothesis selection was acting at these concentrations. Worryingly, cefotaxime exposure enriched for well-known opportunistic Gram negative pathogens including *P. aeruginosa*, which readily infects immunocompromised individuals such as cystic fibrosis patients; and *A. baumanii*, which is most commonly associated with hospital-acquired infections [42]. Enrichment these opportunistic pathogens and human and gut commensals such as *B. fragilis* and *E. faecalis* under cefotaxime exposure may be due to intrinsic resistance which is likely to result in enrichment within communities including susceptible strains.

In summary, our findings develop understanding of selection for antibiotic resistance by showing that the strength of selection for clinically important resistance genes in a community context can be equal across a very large antibiotic concentration gradient; in other words, we show the strength of selection within a given selective space may be constant. Therefore, selection pressure below the MIC of susceptible bacteria may be as strong as selection between the MICs of susceptible and resistant bacteria (traditional selective window). We hypothesise that in our study, this observation is due to the community-wide benefit provided by resistant bacteria harbouring degradative resistance mechanisms. Marked increases in common Gram negative, opportunistic pathogens and co-selection for resistance to other antibiotic classes raises concerns about selection and co-selection for clinically relevant genes (such as *bla*_CTX-M_) in pathogenic hosts occurring in a wide range of ecological compartments.

## MATERIALS AND METHODS

**Complex community collection, storage and preparation.** Domestic sewage influent from a waste water treatment plant serving a small town was collected in October 2015. The treatment plant serves a population of 43 000. Single use aliquots were mixed in a 1:1 ratio with 20 % glycerol, vortexed and stored at −80 °C. Before use, samples were spun down at 21,100 *g* for 10 minutes, the supernatant removed, and the pellet resuspended twice in equal volume of 0.85% NaCl to prevent nutrient/chemical carry over.

A pilot experiment (data not shown) was conducted to determine the appropriate density of the complex community inoculum by comparing growth over 24hours of different dilutions of inoculum at a range of antibiotic concentrations in a 96 well plate. Following results from the pilot experiment, the 10 % volume/volume waste water inoculum was used for all further experiments using the complex community on the basis it produced the tightest replicates at all the time points and the most reliable growth phases over 24 hours.

**Selection experiments.** Iso-sensitest broth (Sigma) was inoculated with 10 % volume/volume of untreated waste water. This was separated into 30 ml aliquots and appropriate amounts of cefotaxime stock solution were added. Cefotaxime (Molekular) stocks were prepared in autoclaved and filtered (0.22 μM) deionised water.

These 30ml aliquots were further separated into 5ml aliquots, with 5 replicates for each of the cefotaxime assay concentrations: 2 mg/L, 1 mg/L and500, 250, 125, 62.5, 31.25, 15.625 and 0 μg/L for the first experiment; and at 128, 64, 32, 16, 8, 4, 2 and 0 mg/L for the second experiment.

All replicates were immediately sampled for the day 0 sampling time point: 2 × 1ml of each replicate for each treatment was spun down at 21 100 ×g for 3 minutes, the supernatant removed and pellet resuspended in 1000 μl 20% glycerol followed by storage at −80 °C. All other samples for DNA extraction were taken following each overnight incubation at 37 °C, 180 rpm shaking, as above.

After each incubation, 50 μl of each microcosm was inoculated into 5ml fresh medium with fresh antibiotic, and samples taken as above for a total of 8 days. Remaining cell suspensions were spun down and stored as above at the end of each experiment.

**Metagenome analyses.** Three replicates were chosen at random from the no antibiotic, 125 μg/L, 500 μg/L and 2 mg/L treatment to undergo shotgun metagenomic sequencing on the MiSeq2 v2 platform at University of Exeter Sequencing Service (ESS).

DNA was extracted from 1 ml of frozen overnight culture using the MoBio extraction kit according to manufacturer’s instructions. DNA was cleaned and concentrated using Ampure™ beads. Firstly, 2 μl of 20 mg/ml RNAse A (Qiagen) was added to 50 μl DNA and incubated for 10 minutes at 37 °C. 50 μl of Ampure™ beads were mixed with the DNA/RNAse solution and mixed gently by pipetting and incubated at room temperature for 5 - 10 minutes. Following pulse centrifugation to collect droplets, tubes were placed on a magnetic stand and left until all beads had precipitated to the side of the tube. Supernatant was removed and beads were washed two times with 300 μl freshly prepared 80 % ethanol. Beads were air dried briefly (1 - 2 minutes), resuspended in 10μl 10mM Tris-HCL and then incubated for another 10 minutes at 50 °C. Following pulse centrifugation and bead precipitation, DNA was transferred into a fresh tube and stored at −20 °C until library preparation and sequencing.

The 12 Nextera Library preparations, quality control, sequencing and primary sequencing analysis (including trimming reads of the barcodes) was performed by ESS. Data was then run through the “online analysis pipeline for antibiotic resistance genes detection from metagenomic data using an integrated structured antibiotic resistance gene database”, the ARGs-OAP [28]. This provides the abundance of different resistance gene classes and subtypes within these groups normalised by parts per million, 16S rRNA copy number, and cell number. For all subsequent analysis, data normalised by 16S rRNA copy number was used to be in accordance with the qPCR data generated. Heatmaps for resistance gene abundance were generated using pandas [43], matplotlib [44] and seaborn [45] Python packages for resistance gene class and beta-lactam resistance gene subtype, for averages of these three replicates.

16S rRNA sequences were extracted from the Illumina sequencing data as follows: first, forward and reverse reads were adaptor trimmed using Skewer [46] in paired end mode. Fastqc [47] and Multiqc [48] verified successful adaptor removal and that sequences were of acceptable quality before paired end reads were combined with FLASH version 2 [49] with the maximum overlap set to 300. 16S rRNA sequences were extracted and assigned to bacterial species using Metaphlan2 [27]. The resulting heatmap of species relative abundance was generated using HClust2 using Bray Curtis distance measurements between samples and features (species) [50]; and overall species abundance across treatments was visualised with GraPhlan [51]. Linear Discriminant Analyses (LDA) effect size (LEfSe) analyses was performed and visualised with LEfSe [52] to identify which species, if any, were enriched by a particular cefotaxime treatment.

**qPCR analyses.** Frozen samples/were thawed and DNA extracted using the MBio UltraClean DNA extraction Kit according to instructions. DNA was diluted 5 – 10x before use. The qPCR conditions were optimised using primers from previous studies [53, 54]: 10 μl Brilliant qPCR SYBR Green master mix, 2 μl primer pair (5 μM each of forward and reverse primers for 16S rRNA, and 9 μM each of forward and reverse CTX-M primers), 0.2 μl BSA (20 mg/ml), 0.4 μl ROX reference dye (20 μM), 5 μl diluted DNA template and filtered, sterilised water to a total volume of 20 μl. The qPCR programme for all reactions was 95 °C for 3 minutes, followed by 40 cycles of 95 °C for 10 seconds and 60 °C for 30 seconds. *bla*_CTX-M_ copy number was divided by 16S rRNA copy number to determine *bla*_CTX-M_ gene/16S rRNA, a proxy for *bla*_CTX-M_ prevalence.

gBlock synthetic genes (IDTDNA - see Table 1) were used as qPCR standards; these were resuspended in TE buffer according to the manufacturer’s instructions and were stored at −80 °C.

**Table 1.**
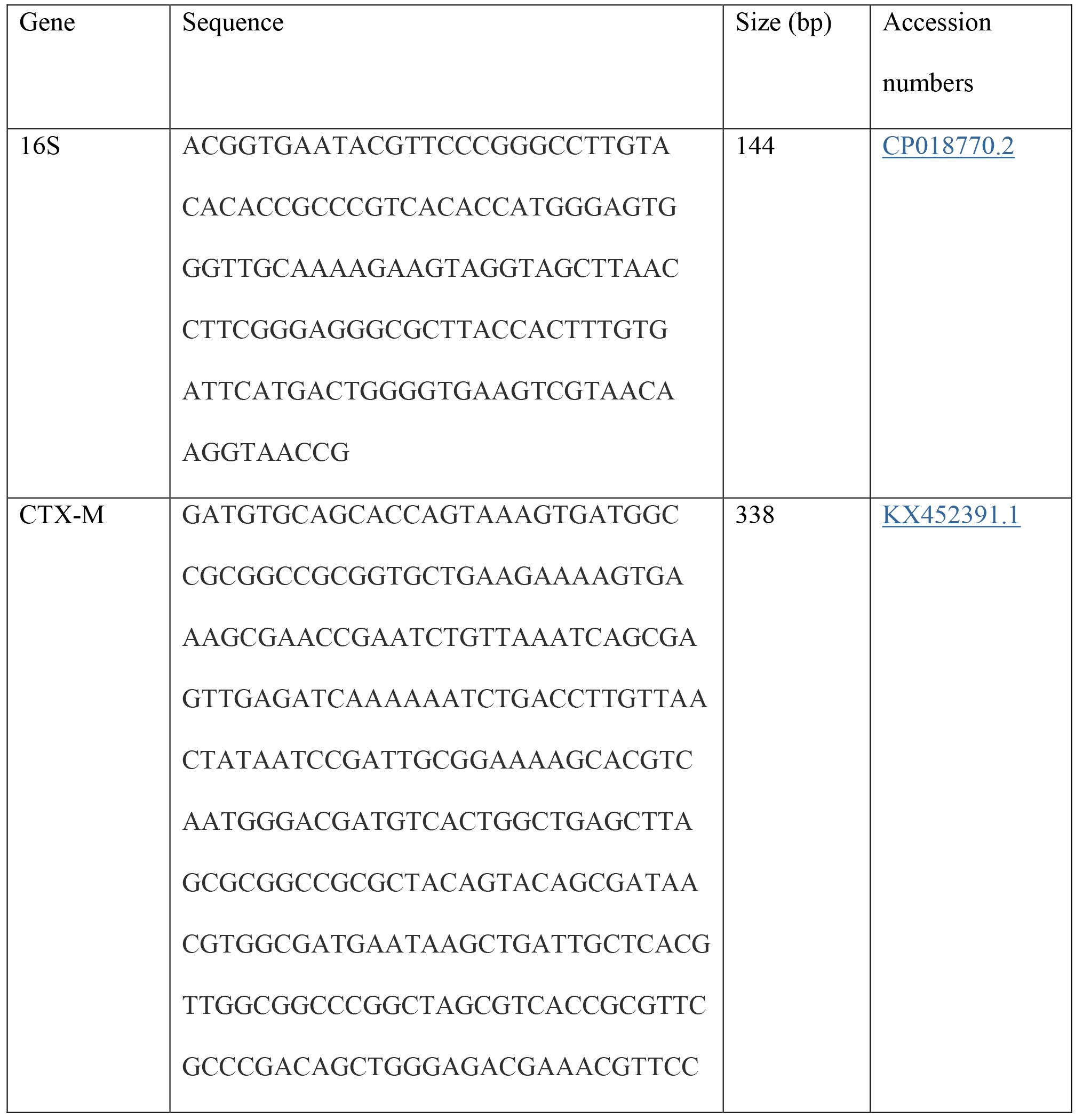
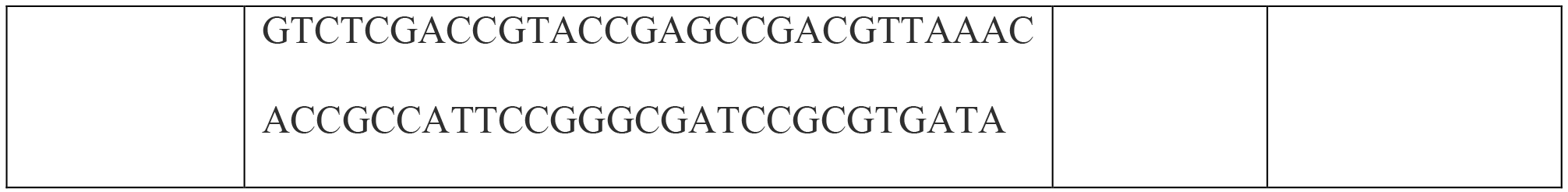
List of synthetic gene blocks used in the study, their DNA sequence, size in bp, and accession numbers used to for design.

Standards were 10 × serially diluted in TE buffer and stored at −20 °C before use. Every PCR plate was always run with 5 serial dilutions of standards in duplicate (and a duplicate negative control). Provided the efficiency for the reaction was between 90 *%* and 110 %, the average CT’s for the duplicate technical replicates for each sample was used to calculate the copy number based on a ‘gold standard’ standard series, where the DNA concentration had been quantified by QuBit and the copy number per μl quantified immediately prior to cycling.

**Data analyses.** All statistics were performed in R Studio [55]. Selection coefficients were used to determine MSC, with the following equation, as previously[3]: [ln(R(t)/R(0))]/[t], where R = resistance prevalence, t = time in days and R(0) = resistance prevalence at time zero. The MSC is estimated by the line of best fit X-axis intersect. Figures were generated with various python packages [43–45].

**Chemical extraction/analyses.** Complex community microcosms were sampled at the Day 0 time point and after the first 24 hours. Antibiotic stocks were also quantified.

The extraction procedure was as follows: 400 μl of culture was mixed with 400 μl HiPerChromosolv Acetonitrile in a 2 ml 96 well plate, and spun at 3500 rpm for 30 minutes. 100 pl of this supernatant was then mixed with 900 μl of 1:4 Acetonitrile to HPLC-grade water in a fresh plate, and stored in the fridge. Antibiotic stocks were diluted to a final concentration of 100 ng/L in 1:4 Acetonitrile. Extractions were kept at 4 °C until processing.

Each concentration from the evolution experiment had a minimum of two chemical replicates from at least two of the biological replicates. Stocks were single replicate only.

Chemical quantification was performed at the University of Exeter Streatham Campus by Maciek Trnzadel and Malcolm Hetherdige, co-funded by AstraZeneca Global SHE and the University of Exeter.

**Antibiotic degradation experiment.** Washed, untreated waste water was diluted in 25ml Iso-sensitest broth aliquots spiked with cefotaxime concentrations of 0, 15.625, 31.25, 62.5, 125, 250, 500 μg/L or 2 mg/L. A sterile control at the same concentrations was also prepared. These were incubated at 37 °C, 180 rpm shaking in between sampling. Chemical extractions (see below) and destructive sampling for OD readings were performed at time 0, then every 3 hours for 24 hours. OD measurements were carried out in a spectrophotometer (Jenway, UK) at the same time points at 600 nm. Any OD readings with a value greater than 1 were diluted 10 × in Iso-sensitest broth and then re-measured.

**Supernatant experiment.** *E.coli* strains J53 and NCTC 13451 were grown overnight at 37 °C, shaking at 180 rpm in Iso-sensitest broth (supplemented with 2 mg/L cefotaxime for NCTC 13451). This concentration was chosen on the basis it was greater than the J53 MIC (> 250 μg/L and <500 μg/L, determined by microdilution assay [56]), that it would be fully degraded in a beta-lactamase producing community (according to the overnight degradation experiment); and because it is the clinical breakpoint concentration for *Enterobacteriaceae* [23]. The supernatants from both overnight cultures were spun down at 21 000 *x g* for 2 minutes twice, and then filtered through a 0.22 μM filter. J53 was inoculated at a starting optical density (600nm) of 0.01 into fresh Iso-sensitest broth amended with 2 mg/L and J53 or NCTC 13451 supernatant (12.5% volume/volume). Controls included a blank control (to check general aseptic technique), broth with each supernatant (to verify the supernatant was sterile); and J53 in broth both with and without antibiotic (to deduce effects of nutrient dilution).

## ACKNOWLEDGEMENTS

Aimee Murray was supported by a BBSRC/AZ CASE Studentship, BB/L502509/1. Lihong Zhang was supported Natural Environment Research Council grant NE/M01133X/1. Chemical extraction and quantification protocols, and chemical quantification were performed by Maciej Trznadel and Malcolm Hetheridge, University of Exeter and were funded by AstraZeneca,

## SUPPLEMENTARY LEGENDS

Figure S1. Heatmap showing relative abundance of all detected species using Bray Curtis distance measurements for treatment (x axis) and species (y axis) for each cefotaxime treatment. ‘C0’, ‘C4’, ‘C6’ and ‘C8’ correspond to 0, 125, 500 and 2000 μg/L cefotaxime respectively. The number after the concentration denotes the biological replicate number (1 – 5), chosen randomly for sequencing at day 8 of the experiment.

Figure S2. Linear Discriminant Analyses (LDA) effect size (LEfSe) analyses of statistically significant species associated with different cefotaxime treatments. Negative LDA scores (red) show species enriched in the no antibiotic treatment, and positive LDA scores (green) show species enriched in the 2000 μg/L cefotaxime treatment.

Figure S3. Heatmap showing average (biological replicate n=3) resistance gene relative abundance (resistance gene number normalised with 16S rRNA copy number), following 8 days culture with cefotaxime. “Macrolide-Linco-Strept” = Macrolide, Lincosamide and Streptogramin resistance.

Figure S4. MSC (cefotaxime concentration at the x-axis intercept) determination using average (n=5) selection coefficients (natural log of *bla*_CTX-M_ prevalence over 8 days, *bla*_CTX-M_ prevalence=*bla*_CTX-M_ copy number/16S rRNA copy number, qPCR technical replicate n=2). Shown with standard error bars (of biological replicates) and polynomial (order 2) line of best fit.

Figure S5. Average (biological replicate n=5, technical qPCR replicate of each biological replicate n=2) bla_CTX-M_ prevalence (bla_CTX-M_ copy number/16S rRNA copy number) number at day 0 and following 1, 4 and 8 days of cefotaxime exposure. Shown with standard error bars (of biological replicates).

Figure S6. Average (biological replicate n=5, technical qPCR replicate of each biological replicate n=2) bla_CTX-M_ copy number following 8 days cefotaxime exposure in the higher concentration experiment. Shown with standard error bars (of biological replicates).

Figure S7. Average (biological replicate n=5, technical qPCR replicate of each biological replicate n=2) 16S rRNA copy number following 8 days cefotaxime exposure, in the higher concentration experiment. Shown with standard error bars (of biological replicates).

Figure S8. Growth (optical density (600nm)) of the complex community over time during the 24 hour degradation experiment. Single replicate only.

Table S1. Nominal (expected) and average (biological replicate n=3, technical replicate of each n=2) measured cefotaxime concentrations as determined by LC-MS at the beginning (time 0) of the selection experiment, and after 24 hours culture at 180rpm, 37°C in the presence of the complex community. Also shown are the cefotaxime stocks (‘1’ and ‘2’) used in the experiment.

